# Microbial Analysis and Sanitization of Hydroponic Farming Facilities in Singapore

**DOI:** 10.1101/2024.04.08.588599

**Authors:** Cliff An Ting Tham, Ye Htut Zwe, Michelle Mei Zhen Ten, Geraldine Shang Ya Ng, Jillinda Yi Ling Toh, Bee Ling Poh, Weibiao Zhou, Dan Li

**Author notes:** Corresponding author: Dr. Dan Li.

## Abstract

This study performed microbial analysis of nutrient film technique (NFT) hydroponic systems at three indoor farms in Singapore. To justify the necessity to sanitize the hydroponic systems, strong biofilm-forming bacteria were isolated from the facility and investigated with their influence on *Salmonella* colonizing on polyvinyl chloride (PVC) coupons in hydroponic nutrient solutions. Last, sanitization solutions were evaluated with both laboratory-scale and field-scale tests. As a result, the microbiome composition in NFT systems was found to be highly farm-specific. Strong biofilm formers *Corynebacterium tuberculostearicum* C2 and *Pseudoxanthomonas mexicana* C3 were found to facilitate the attachment and colonization of *Salmonella* on PVC coupons. When forming dual-species biofilms, the presence of C2 and C3 also significantly promoted the growth of *Salmonella* (*P* < 0.05). Sodium hypochlorite (NaOCl) exhibited superior efficacy in biofilm removal compared to hydrogen peroxide (H_2_O_2_) and sodium percarbonate (SPC). NaOCl at 50 ppm reduced C2 and C3 counts to < 1 log CFU/cm^2^ within 12 h, whereas neither 3% H_2_O_2_ nor 1% SPC achieved such an effect. In operational hydroponic systems, the concentration of NaOCl needed to achieve biofilm elimination increased to 500 ppm, likely due to the presence of organic matter accumulated during the crop cultivation and the higher persistence of the naturally formed multispecies biofilms. The sanitization (500 ppm NaOCl for 12 h) did not impede subsequent plant growth but chlorination by-product chlorate was detected with high levels from the hydroponic solution and plants in the sanitized systems without rinsing.

**IMPORTANCE:** This study’s significance lies first in its elucidation of the necessity to sanitize the hydroponic farming systems. The microbiome in hydroponic systems, although most of the times non-pathogenic, might serve as a hotbed for pathogens’ colonization and thus pose a higher risk for food safety. We thus explored sanitization solutions with both laboratory-scale and field-scale tests. Of the three tested sanitizers, NaOCl was the most effective and economical option, whereas one must note the vital importance of rinsing the hydroponic systems after sanitization with NaOCl.

## INTRODUCTION

Urbanization poses significant challenges to food security worldwide, particularly in densely populated areas where arable land is scarce (1). Singapore, emblematic of such urban centers, grapples with the conundrum of sustaining its food supply amidst limited land availability for traditional agriculture. With only 1% of its land designated for farming due to competing land uses, Singapore relies heavily on food imports to meet over 90% of its nutritional needs (2). This dependency exposes the nation to vulnerabilities stemming from disruptions in global food supply chains. To address this pressing issue, Singapore launched the “30 by 30” initiative in 2019, aiming to produce 30% of its nutritional requirements locally by 2030 (3). Central to this initiative is the exploration of innovative agricultural practices capable of maximizing food production within the constraints of urban spaces. Among these practices, hydroponic farming has emerged as a promising solution, offering the potential to cultivate vegetables sustainably in indoor environments with minimal land requirements (4).

However, indoor growing with precise control of environmental variables such as temperature and carbon dioxide level does not automatically ensure that the produce is exempt from food safety issues. Even with stringent controls, undesired microorganisms could still be introduced into the indoor growing systems from multiple sources including water, fertilizer, growth substrate, seeds/seedlings, and pests, as well as contamination from human manipulation. Unlike traditional soil-based farming, hydroponic systems provide a conducive environment for microbial biofilm formation on various surfaces, including those in nutrient film technique (NFT) systems commonly employed in indoor farms (5–6). Our previous study demonstrated that once the pathogenic bacteria *Salmonella* was introduced, it could rapidly spread to the whole hydroponic system likely by the nutrient solution circulation (7).

*Salmonella* is a leading cause of bacterial foodborne illnesses and outbreaks worldwide and is heavily correlated with the consumption of fresh produce supported by numerous pieces of evidence (8–11). In 2021, eight varieties of hydroponic greenhouse-grown leafy green vegetables from BrightFarms brand have been linked to a *Salmonella* outbreak. Eleven people became ill, two were hospitalized, and there were likely more unreported cases of illness (estimated to be 30 times of the reported cases) during this outbreak (12). During a screening study conducted by our group (13), multiple *Salmonella* strains belonging to seven different serovars were isolated from both the nutrient solution and the lettuce crops from local farms in Singapore, suggesting the need for local farms to improve their sanitation and hygiene practices.

Addressing microbial contamination in hydroponic systems necessitates the development and implementation of robust cleaning and sanitization protocols, which must consider not only their efficacy but also their practicality, cost-effectiveness, and potential impact on plant health and system sustainability. Balancing these factors is crucial for ensuring the viability and resilience of hydroponic farming practices in urban settings like Singapore. This study aimed to first accumulate baseline knowledge of the microbial loads and composition in operational indoor hydroponic farms in Singapore. By investigating the biofilm-forming capabilities of the bacterial isolates from the hydroponic farms and the interactions of the strong biofilm formers with *Salmonella*, one of the most representative pathogens, we intended to elucidate the necessity of sanitizing the hydroponic farming systems. Accordingly, our work tested and proposed effective sanitation measures to safeguard the food safety of indoor hydroponic farming systems.

## RESULTS AND DISCUSSION

### Microbial analysis of multi-tier NFT hydroponic systems

The microbial analysis was conducted by collecting samples from the NFT hydroponic systems in operation at three commercial indoor farms in Singapore. Ten surface swabbing samples were collected from each of the seven components of the NFT systems as illustrated in Figure 1b. Interestingly, it was found that at all three farms, the nutrient tank interior surfaces and main discharge points leading to the nutrient tanks always exhibited significantly higher microbial counts (7.3-7.5 log CFU/cm^2^) than the other tested parts (*P* < 0.05; Figure 1a).

**Figure 1.**
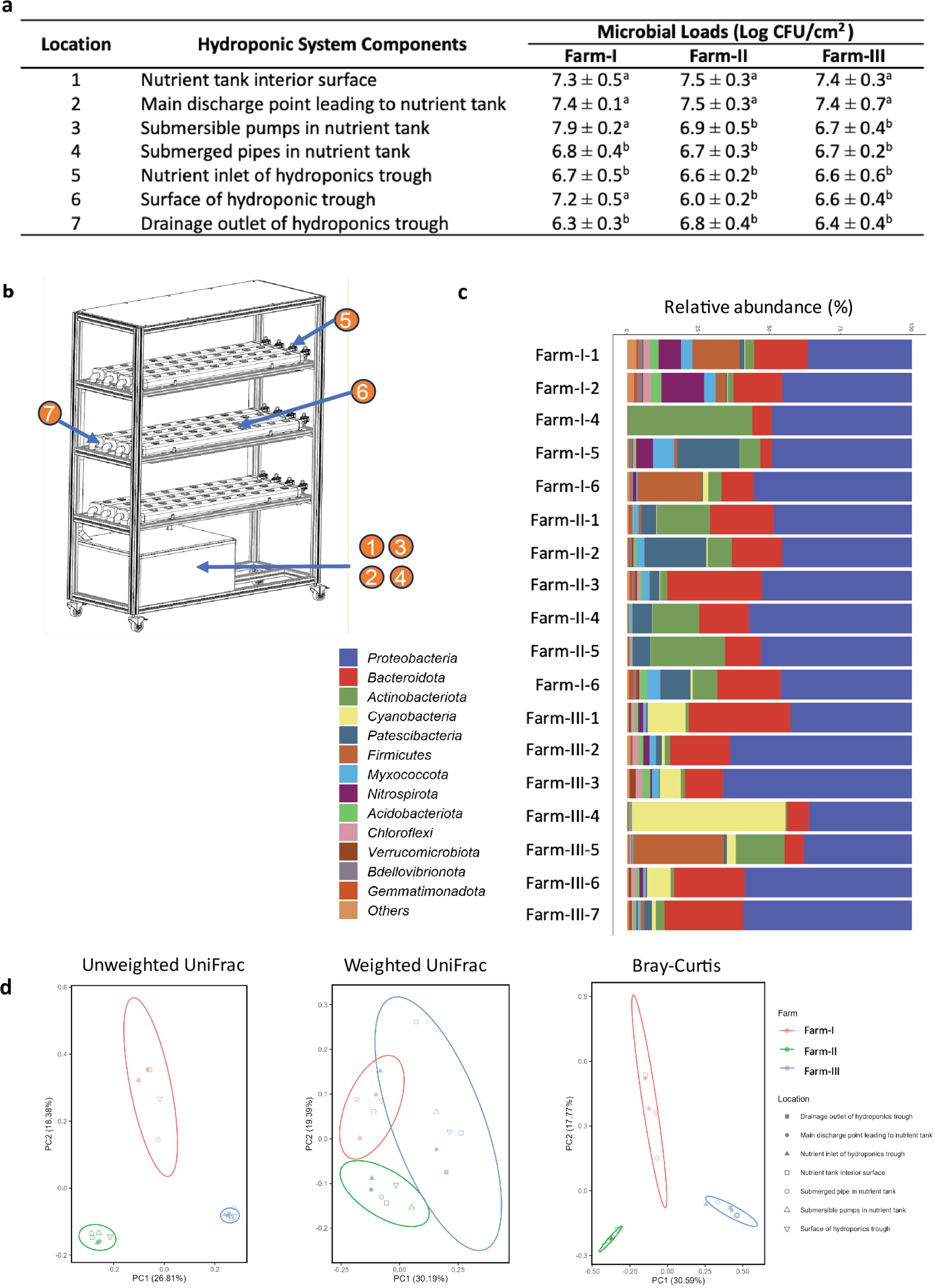
The key microbial analysis results of NFT hydroponic systems at three commercial indoor farms in Singapore. **(a)** The microbial loads in different components of NFT systems. Values denoted with different letters are significantly different (*P* < 0.05). For each data point, 10 samples were collected from each farm (n = 30). **(b)** The illustration of the NFT systems. **(c)** The identified bacteria phyla by 16S rDNA sequencing (> 1%) from selected samples. **(d)** Microbiome compositional difference (beta-diversity) based on the bacterial composition analysis. Unweighted UniFrac (left, presence of taxa), weighted UniFrac (middle, presence and relative abundance of taxa), and Bray-Curtis (right, presence and relative abundance of taxa) distance are visualized by PCoA plots.

The bacteria composition of the swabbed surfaces was identified by 16S rDNA sequencing. At phylum-level, Proteobacteria was the most abundant group from all the tested samples (36.0 – 63.8%) except for the submerged pipes in the nutrient tank at Farm-I. Next, Bacteroidota was also identified from all of the samples with major relative abundances (6.8 – 35.8%) (Figure 1c). The microbiome composition became very diverse at the genus-level (Figure S1), making it difficult to identify the commonly shared genera from all the samples. Instead, the genera composition tended to be more farm-specific. At Farm-I, unclassified *Comamonadaceae, Nitrospira, Bacillus*, unclassified *Rhizobiaceae, Aeromonas* were identified with higher than 1 % relative abundances from more than half of the samples. More genera were identified when the same criteria (higher than 1% relative abundances from more than half samples) were applied to the samples from Farm-II, including unclassified *Comamonadaceae*, *Sphingobium*, Saccharimonadaceae TM7a, *Pseudomonas, Hyphomicrobium, Terrimonas,* unclassified *Chitinophagaceae, Panacagrimonas, Microbacterium, Thermomonas, Saccharimonadales, Luteimonas,* and *Allorhizobium-Neorhizobium-Pararhizobium-Rhizobium.* Diverse genera were also identified for samples from Farm-III being mostly different from samples of Farm-I and Farm-II. Genera with higher than 1% relative abundances in more than half of the samples at Farm-III included *Hydrogenophaga, Leptolyngbya* PCC-6306, Unclassified *Saprospiraceae*, unclassified *Comamonadaceae*, *Cyanobacteria, Haliscomenobacter, Rhodobacter, Hyphomicrobium, Porphyrobacter, Hyphomonadaceae* SWB02, *Sediminibacterium,* and *Ferrovibrio.* Indeed, the beta-diversity analysis over the amplicon sequence variants (ASVs) also revealed the similarities in the microbiome shared by samples from the same farm. Clear clusters for samples from the three farms were identified from the principal coordinate analysis (PCoA) plots based on weighted and unweighted UniFrac distance, as well as count-based Bray-Curtis distance (Figure 1d). Since the three farms were all indoor hydroponic farms using NFT systems with polyvinyl chloride (PVC) materials, using tap water to formulate the hydroponic solutions, and growing Brassica leafy greens, the most probable reasons causing this distinctive farm-specific microbiome composition could be the personnel and other environmental factors.

In addition, we have also performed 18S rDNA sequencing to identify the fungi and algae from the surface samples. Microalgae belonging to Chlorophyta, Ochrophyta, Diatomea, and fungi belonging to Ascomycota, Cryptomycota, LKM15 fungi were identified from most of the samples from all three farms (Figure S2). Specifically, the green algae Chlorophyta was found with high relative abundances in most of the samples, especially from Farm-III samples (> 90%, Figure S2). Very recently, Lipsman et al. (14) revealed that a shared algal-bacterial extracellular matrix could promote algal-bacterial aggregation in an environmentally relevant model system. In their study, algal exudates played a pivotal role in promoting bacterial colonization, stimulating bacterial exopolysaccharide production, and facilitating a joint formation of algal-bacterial extracellular matrix. So far, it remains largely unknown over the influence of green algae on food safety of the hydroponic systems, which is thus worthwhile for exploration in the future. In this study, the following parts were still mainly focused on the bacteria. The influence of the background microbiome on the microbial safety of hydroponic systems was investigated by evaluating the interactions of the bacterial biofilm formers with *Salmonella*, one of the most representative foodborne pathogens. The evaluation of sanitization efficacy was evaluated by plate counts on plate count agar, a non-selective agar mainly assessing total viable bacteria.

### Biofilm former identification

Ten bacteria colonies with various morphology were isolated from the samples from the nutrient tank interior surfaces. Their biofilm-forming abilities were observed by the measurement of biomass (Figure 2a). With the use of the criteria established previously (15), 4 isolates, as well as the *Salmonella* serovar Brunei (a self-isolate from a hydroponic farm in Singapore by our group from a previous study [13]), were attributed as weak biofilm formers, 3 isolates were attributed as moderate biofilm formers, and 3 isolates were attributed as strong biofilm formers. We thus selected isolates C2 and C3, the strongest gram-positive and gram-negative biofilm formers respectively for the following studies. Whole genome sequencing (WGS) analysis identified C2 as *Corynebacterium tuberculostearicum* (Figure 2b) and C3 as *Pseudoxanthomonas mexicana* (Figure 2c).

**Figure 2.**
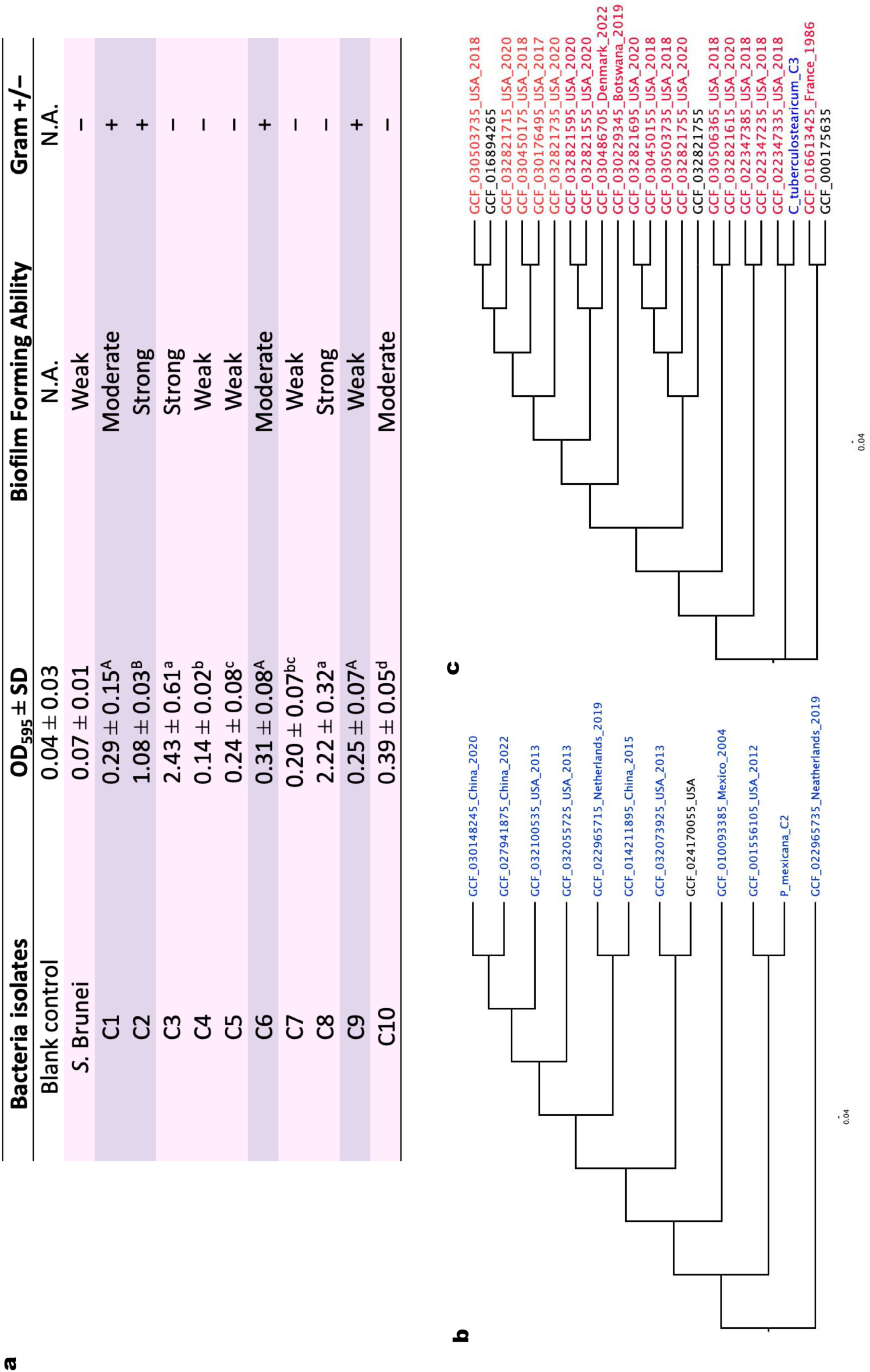
Screening of strong biofilm formers from the hydroponic systems’ interior surfaces. **(a)** Biofilm forming capability and gram staining results of the 10 bacterial isolates. Results are based on the average of triplicates. A-B denote the significance between gram-positive strains, a-d denote the significance between gram-negative strains. **(b)** and **(c)** show the SNP trees based on the maximum likelihood method illustrating the phylogenetic relationship between C2 and C3 with the closely related members of the same genus, respectively. The numbers represent bootstrap values. The red IDs represent human clinical isolates, the ones with blue IDs are environmental isolates, and the strains with black IDs are with unknown origins.

### Effect of *C. tuberculostearicum* C2 and *P. mexicana* C3 biofilms on the colonization of *Salmonella* on PVC coupons

The *S.* Brunei strain used in this study showed weak biofilm-forming capability as shown in Figure 2a. When we dipped PVC coupons in a 5 Log CFU/mL *Salmonella* suspension and rinsed the coupons with sterile deionized water, no *Salmonella* was enumerated from the PVC coupons (< 1 Log CFU/cm^2^) after up to 120 h incubation of the coupons in hydroponic nutrient solution (Figure 3a). In contrast, on PVC coupons with pre-formed *C. tuberculostearicum* C2 and *P. mexicana* C3 biofilms, *Salmonella* was detected at 2.2 ± 0.3 and 1.9 ± 0.2 Log CFU/cm^2^ after water rinsing, and the *Salmonella* counts continued to increase along with the incubation of the coupons in hydroponic nutrient solutions up to 3.3 ± 0.2 and 2.8 ± 0.1 Log CFU/cm^2^ respectively after 120 h (Figure 3a). *C. tuberculostearicum* has been reported as a ubiquitous bacterium that colonizes human skin (16). The occurrence of this bacteria in the hydroponic facility might be due to human manipulation. Although there have been occasional reports of human infection caused by this bacteria species (17), no evidence could be found of any food-related infection or risks. *P. mexicana* is an environmental non-pathogenic bacterium (18). Our results suggest that background microbiomes such as *C. tuberculostearicum* and *P. mexicana,* although being mostly non-pathogenic, could enhance the retaining of foodborne pathogens such as *Salmonella* after being introduced in the hydroponic circulating systems, which might not be able to colonize easily in a cleaner system.

**Figure 3.**
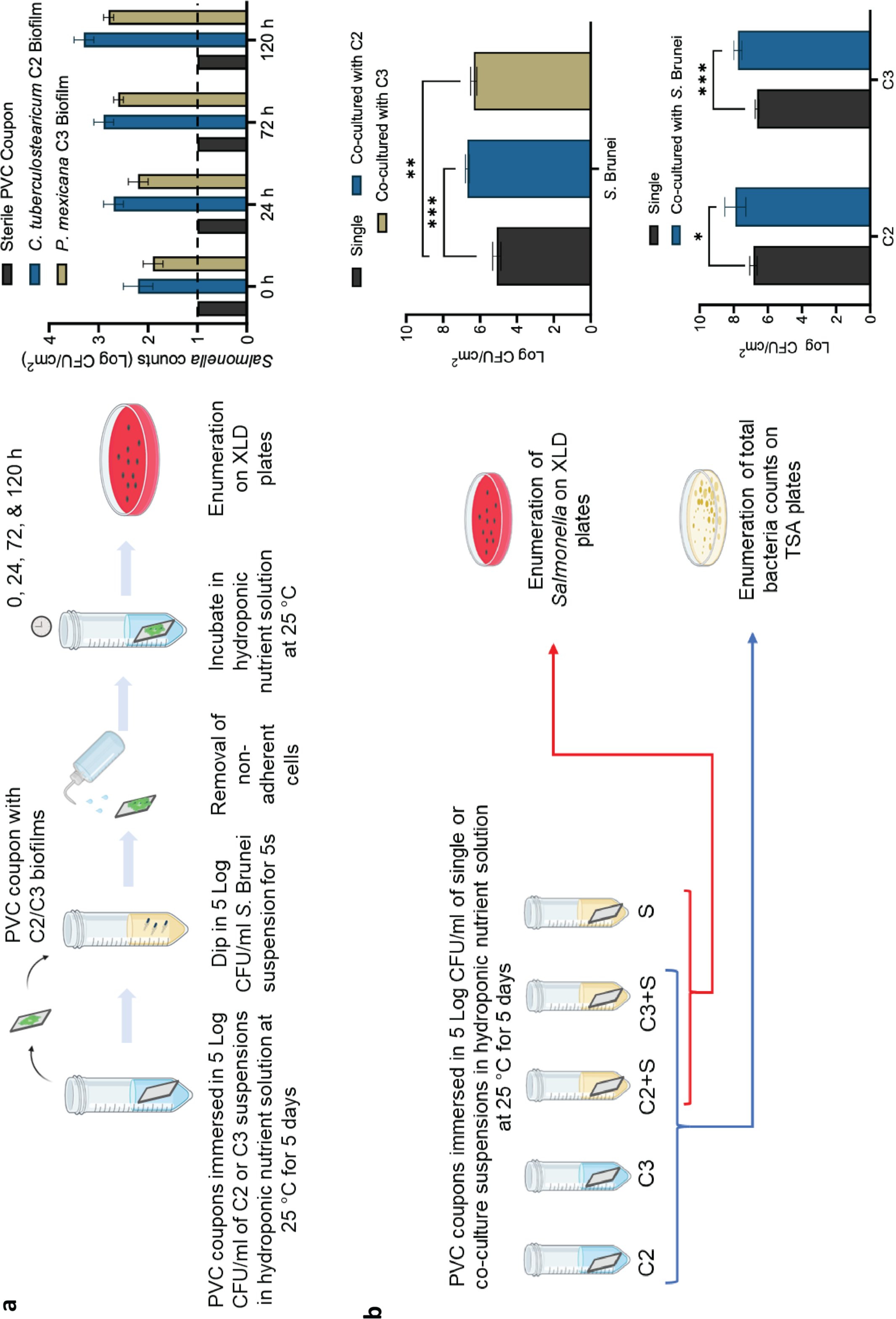
Experimental set-ups and results showing the effectiveness of *C. tuberculostearicum* C2 and *P. mexicana* C3 on the colonization of *S.* Brunei on PVC coupons. **(a)** Adherence and colonization of *Salmonella* on pre-existing *C. tuberculostearicum* C2 and *P. mexicana* C3 biofilms. Dotted line represents the limit of detection. **(b)** *Salmonella* (on XLD plates) and total bacteria counts (on TSA plates) from single– and dual-species biofilms. Each value is the average of triplicates from three independent experiments. The error bars indicate the standard deviations (SD). *, ** and *** denote *P* < 0.05, 0.01, and 0.001, respectively.

### Effect of *C. tuberculostearicum* C2 and *P. mexicana* C3 on the growth of *Salmonella* in dual-species biofilms

A different set-up was designed and demonstrated *C. tuberculostearicum* C2 and *P. mexicana* C3 could not only assist the adherence of *Salmonella* on surfaces but also could promote the growth of *Salmonella* to higher levels when mixed in dual-species biofilms. As shown in Figure 3b, the *Salmonella* counts were significantly higher in dual-species biofilms co-cultured with *C. tuberculostearicum* C2 (6.7 ± 0.1 Log CFU/cm^2^; *P* = 0.0002) and with *P. mexicana* C3 (6.3 ± 0.2 Log CFU/cm^2^; *P* = 0.0018) compared to its single species-biofilms (5.1 ± 0.2 Log CFU/cm^2^). Due to a lack of selective culture media for *C. tuberculostearicum* and *P. mexicana,* it was impossible to make a direct comparison of the C2 and C3 counts in single– and dual-species biofilms. However, we did observe significantly higher total bacterial counts in the dual-species biofilms (7.9 ± 0.5 and 7.8 ± 0.2 Log CFU/cm^2^) than in the C2 and C3 single-species biofilms (6.9 ± 0.2 and 6.6 ± 0.1 Log CFU/cm^2^; *P* < 0.05; Figure 3b), suggesting a symbiotic interaction between *Salmonella* with *C. tuberculostearicum* and *P. mexicana*.

Such symbiotic effects between different bacterial species in biofilms have been observed in previous studies such as the co-culture biofilm assays involving *Staphylococcus aureus* with *Pseudomonas aeruginosa* (19), and *Staphylococcus epidermidis* with *Candida albicans* (20). However, not all bacterial interactions within mixed-species biofilms are synergistic. For instance, in one of our previous studies, we found that a strain of *Chromobacterium haemolyticum* isolated from hydroponic nutrient solution exhibited highly antagonistic properties, eliminating *Salmonella*, *Escherichia coli*, *Listeria monocytogenes*, and *S. aureus* in co-culture biofilms (21). Nevertheless, barring exceptionally antagonistic strains, our findings again suggest that the presence of resident biofilms in hydroponic systems may increase the risk of the persistence of foodborne pathogens.

### Laboratory-scale test: efficacy of various sanitizers on *C. tuberculostearicum* C2 and *P. mexicana* C3 biofilms

Using *C. tuberculostearicum* C2 and *P. mexicana* C3 biofilms grown on PVC coupons as a representative and robust model, we evaluated the potential of a few commonly used sanitizers, including sodium hypochlorite (NaOCl), hydrogen peroxide (H_2_O_2_), and sodium percarbonate (SPC), being used to sanitize hydroponic systems. The tested concentrations of each sanitizer (50 and 500 ppm NaOCl, 1.5% and 3.0% H_2_O_2_, 1.0% SPC) were selected based on the commonly used concentrations under other application scenarios (22–24). The tested durations (1 h, 6 h, 12 h) were selected based on the practical considerations not to compromise the farming productivity (maximumly overnight sanitization).

As a result, NaOCl exhibited superior efficacy in biofilm removal compared to H_2_O_2_ and SPC. For both *C. tuberculostearicum* C2 and *P. mexicana* C3 biofilms, NaOCl at 500 ppm reduced the bacteria counts to < 1 Log CFU/cm^2^ from the biofilm grown on PVC coupons within 1 h. At a lower concentration of 50 ppm, *C. tuberculostearicum* C2 counts were reduced to < 1 Log CFU/cm^2^ within 12 h, and *P. mexicana* C3 counts were reduced to < 1 Log CFU/cm^2^ within 6 h. In comparison, neither 3% H_2_O_2_ nor 1% SPC achieved such an effect within the tested 12 h (Figure 4).

**Figure 4.**
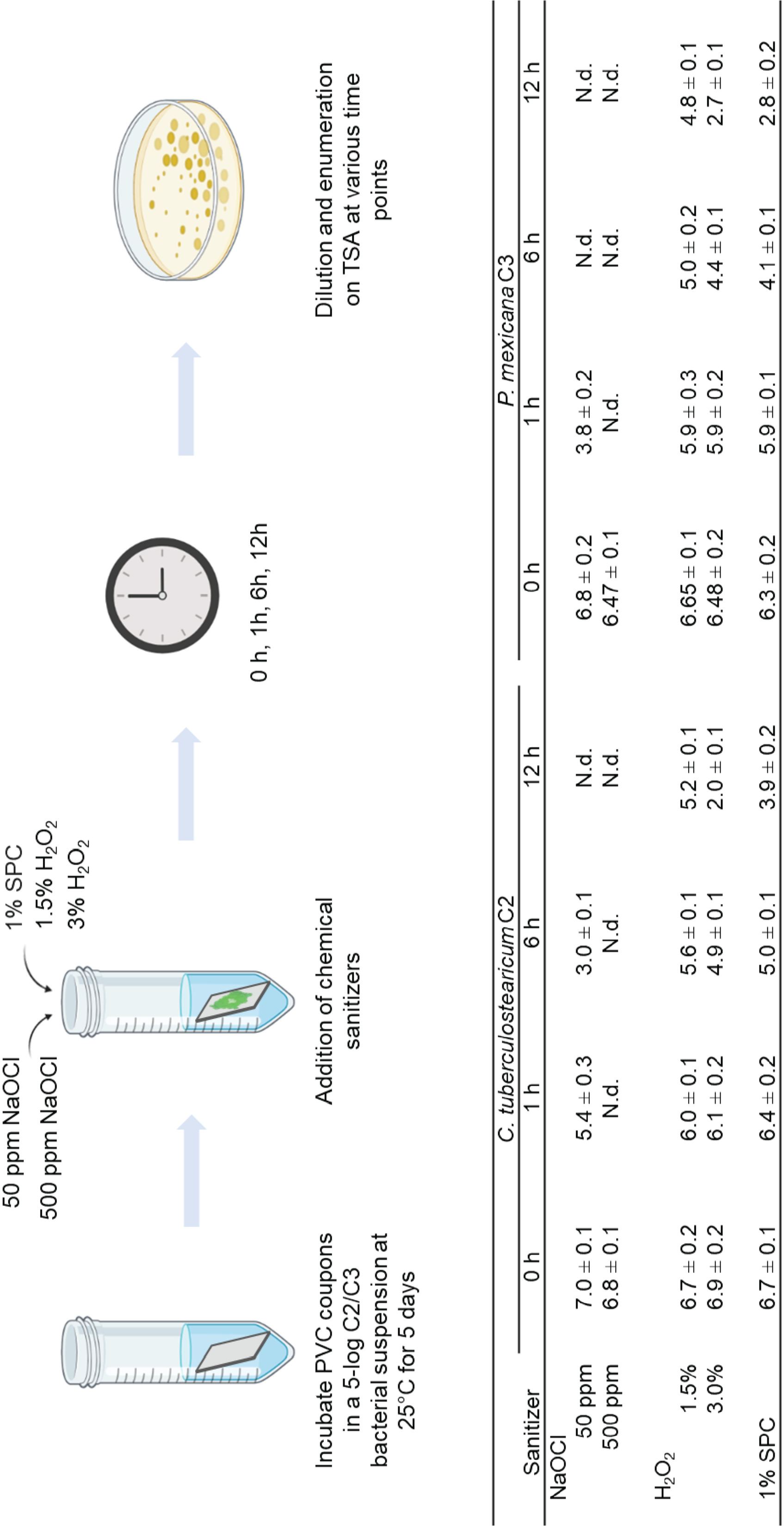
Experimental set-ups and results of sanitization of *C. tuberculostearicum* C2 and *P. mexicana* C3 biofilms grown on PVC coupons. Each value (Log CFU/cm^2^ ± SD) is the average of triplicates from three independent experiments. N.d. denotes not detected, < 1 Log CFU/cm^2^.

### Field-scale test: sanitization of commercial NFT hydroponic systems

The dose of NaOCl and duration required for sanitization of commercial NFT hydroponic systems in the field-scale test increased remarkably in comparison with the laboratory-scale test. As shown in Table 1, the microbial loads on the NFT system surfaces were only reduced to levels < 1 Log CFU/cm^2^ by 500 ppm NaOCl treatment for 12 h. On one hand, the mixed-species biofilms have been reportedly repeatedly with higher resistance toward multiple inactivation treatments (25–27). On the other hand, this was also likely due to the presence of organic matter accumulated during crop cultivation. Organic matters such as protein and carbohydrates are well-known to induce significant interference with the sanitizing effect of NaOCl (28–29). In this study, the concentrated sanitizers were directly added to the hydroponic solution in the NFT systems reaching the defined working concentrations after the crop harvest. Although replacing the hydroponic solution with fresh sanitizers is expected to generate better efficacy by reducing the organic matter loads, one must consider the sustainability and economic factors at the same time when designing the protocol.

**Table 1.**
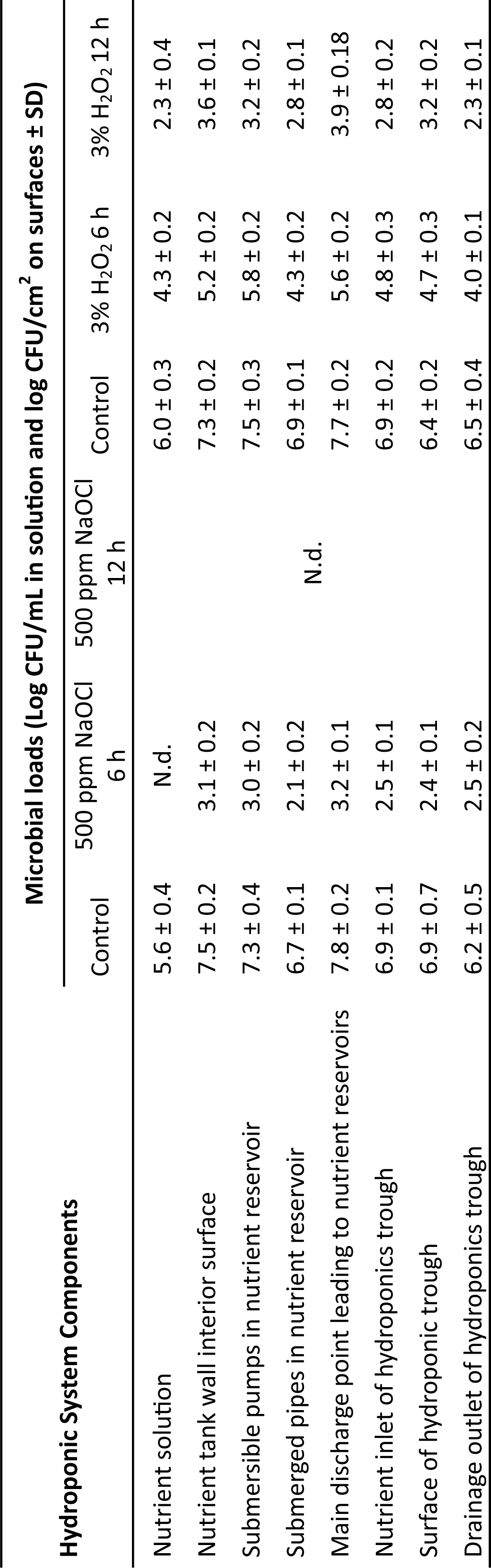
Effcacy of 500 ppm NaOCl and 3% H_2_O_2_ in sanitizing the multi-tier NFT hydroponic systems at three commercial indoor farms in Singapore. For each data point, 10 samples were collected from each farm (n = 30). N.d. denotes not detected, < 1 log CFU/cm^2^ or < 1 log CFU/mL.

Being consistent with the laboratory-scale test, H_2_O_2_ was less effective in eliminating the microbial loads in the NFT systems. Although a time-dependent effect was observed within all tested components resulting in significant reductions (*P* < 0.05), 2.3-3.9 Log CFU/cm^2^ of microbial counts were still obtained after treatment by 3% H_2_O_2_ for 12 h (Table 1). H_2_O_2_ has been gaining high popularity among hydroponic growers due to the free concern over the chemical residuals, leading to many studies evaluating its efficacy (30–34) and commercial companies providing H_2_O_2_-based hydroponic decontamination solutions such as Waterco from Australia. Furthermore, the literature also extensively documented the potential role of H_2_O_2_ as a signaling molecule in abiotic stress response and photosynthesis, contributing to growth performance in plants (35–37) and thus further enhancing the appeal of H_2_O_2_ for hydroponic growers. However, beyond efficacy, economic and practical factors also favor NaOCl. For instance, in Singapore, the cost per liter of 3% H_2_O_2_ exceeds that of 5% NaOCl (household bleach) by approximately 3.5 times. Since our results advocate for diluting commercially available 5% NaOCl by a factor of 100 before application, it further reduces the cost of sanitizing the systems with NaOCl.

### Influence of NaOCl sanitization on the yield and chemical safety of the indoor grown agricultural products

NaOCl, although being effective and economical as a commonly used sanitizer, is associated with significant chemical safety concerns due to the generation of harmful by-products (38–41). For instance, the use of chlorine in fresh-cut produce washing is prohibited in some European Union countries including Germany, Switzerland, the Netherlands, Denmark, and Belgium (42–43).

In this study, following a 12-h treatment with 500 ppm NaOCl, the hydroponic system was drained and replenished with fresh nutrient solution without additional rinsing. Komatsuna (*Brassica rapa* var. *perviridis*) seeds were germinated in this system. As shown in Table 2, after 7 days, the komatsuna seeds germinated in the sanitized system achieved a germination rate of 98%. At the end of the growth circle (Day 35), the harvested crops had fresh weight of 109.0 ± 9.6 g/plant, and 12.5 ± 2.9 leaves/plant. In comparison, seeds germinated in systems without sanitization achieved a germination rate of 99%. At the end of the growth circle (Day 35), the crops had fresh weight of 109.0 ± 9.2 g/plant, and 12.8 ± 2.7 leaves/plant. No significant differences (*P* > 0.05) in fresh weight and number of leaves were observed.

**Table 2.** Germination rate (n = 100, Day 7) and yield (fresh weight and number of leaves, n = 30, Day 35) of komatsuna (*Brassica rapa* var. *perviridis*) grown in nutrient solution sanitization with 500 ppm NaOCl for 12 h.

| In sanitized system |  |  | In unsanitized system |  |  |
| --- | --- | --- | --- | --- | --- |
| Germination rate (%) | Fresh weight (g) | Number of leaves | Germination rate (%) | Fresh weight (g) | Number of leaves |
| 98 | 109.6 ± 9.6 | 13.0 ± 3.0 | 99 | 109.7 ± 9.2 | 13.0 ± 3.0 |

At the end of the growth circle (Day 35), we took the nutrient solution sample in the hydroponic system and analyzed a series of chlorination by-products. As a result, the chlorate presence was at an alarmingly high level (2383 mg/L) while the rest were below detection limits (Table 3a). We thus also analyzed the level of chlorate in the harvested komatsuna and detected a level of 0.054 mg/kg chlorate in the vegetables (Table 3b).

**Table 3.**
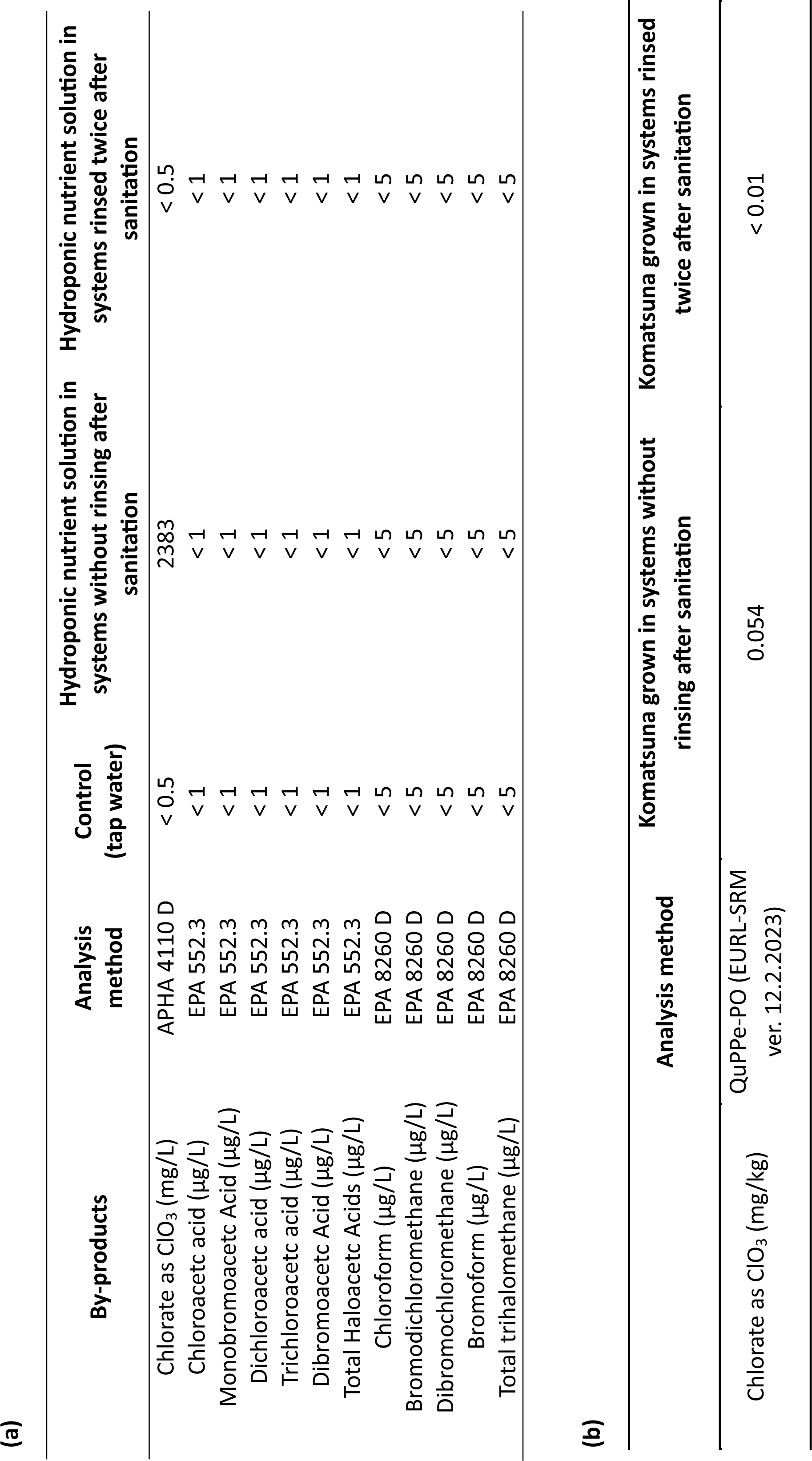
Analysis of chlorination by-products present in the hydroponics nutrient solution in the NFT systems (a) and in ready-to-harvest komatsuna (*Brassica rapa* var. *perviridis*) grown in the NFT systems for 35 days (b).

The European Food Safety Authority (EFSA) has estimated the risks of chlorate in human health (44). The recommended safe intake level for a daily intake of chlorate is 36 µg/kg of body weight per day, as a high intake of chlorate can limit the blood’s ability to absorb oxygen, leading to kidney failure. Over time, chronic exposure to chlorate can inhibit iodine uptake. EFSA has thus also set a tolerable daily intake (TDI) of 3 µg/kg of body weight per day for long-term exposure to chlorate in food. Based on this recommendation, the chlorate intake due to the consumption of vegetables grown in the sanitized hydroponic systems in this study would barely reach the TDI. However, one may not rule out the possibility of chlorate intake from other sources such as drinking water. Therefore, rinsing steps after sanitization became essential to avoid introducing new hazards into the hydroponic-grown agricultural products. In our next trial, the systems were rinsed with tap water twice after sanitization before the growth. The chlorate levels of both the hydroponic solution and the vegetables were reduced to levels below detection limits (< 0.5 mg/L in the hydroponic solution and < 0.01 mg/kg in the vegetables, Table 3a and 3b).

Taken together, our results suggest that omitting the rinsing steps would not diminish the crop yield but the harmful by-products could remain in high levels in the hydroponic solutions and be internalized and accumulated in the edible parts of the crops grown in the systems, which are consequences that could be easily ignored by the growers and consumers, and potentially become a new food safety issue in the long run. It is therefore vitally important to set regulatory guidelines over the technical details of sanitizing hydroponic systems with NaOCl, especially to prevent overdose and to ensure the rinsing steps.

## CONCLUSIONS

Utilizing hydroponics to address food security concerns in a highly urbanized country such as Singapore comes with its own set of unique challenges. Without proper sanitization measures, the closed-loop process heightens the risk of microbial contamination and persistence within the systems. In this study, we demonstrated the high microbial loads as well as the diverse and farm-specific microbiome compositions in the hydroponic systems at three commercial farms in Singapore. Two strong biofilm-forming bacteria isolated from the interior surfaces of the hydroponic systems were able to facilitate the colonization of *Salmonella* on PVC coupons in hydroponic solutions. Last, we compared the efficacy of NaOCl, H_2_O_2_, and SPC in sanitizing the hydroponic systems. NaOCl turned out to be the most effective and economical option, whereas a rinsing step was found to be of vital importance to avoid the harmful chlorination by-products in the final products. This study contributes valuable insights into the management of microbial contamination in hydroponic farming, facilitating the advancement of sustainable agriculture practices in urban environments like Singapore.

## MATERIALS AND METHODS

### Microbial analysis of multi-tier NFT hydroponic systems

Samples were collected from three indoor farms using multi-tier NFT hydroponic systems to grow Brassica leafy vegetables in Singapore between August and September of 2023. All three farms had maintained continuous operation for a period exceeding three months before the sampling. After harvest, the nutrient solution was drained, and seven distinct components of the hydroponic system as listed in Figure 1a and illustrated in Figure 1b were swabbed in 1 cm × 1 cm area with sterile cotton swabs as one sample. Ten samples were swabbed for each component at each farm.

The swabs were submerged in 10 mL of phosphate buffered saline (PBS; Vivantis) in individual 50 mL falcon tubes, transported to our laboratory on ice, and analyzed on the same day. After vortex for 10 s, 1 mL of each 10 mL of the suspension in the falcon tubes was serially diluted in 0.1% peptone water (PW; Oxoid) and enumerated on plate count agar (PCA; Oxoid) after 3 days incubation at 30°C.

### DNA extraction and 16s and 18s rDNA gene sequencing analysis

DNA was extracted from the remnant 9 mL suspensions from the swabs after taking 1 mL for microbial enumeration. Each 9 mL suspension was centrifuged at 9,000 *× g* for 5 min (Centrifuge 5810 R, Eppendorf, Hamburg, Germany). DNA extraction was performed from the pellet using the GeneJET Genomic DNA Purification Kit (Thermo Fisher Scientific, Vilnius, Lithuania) according to the manufacturer’s instructions. The quantity and quality of DNA samples were assessed with BioDrop DUO (BioDrop, Cambridge, United Kingdom), after which the samples were stored at – 80°C for downstream sequencing and analysis.

The 16s and 18s rDNA gene amplification and sequencing were performed at Azenta US Inc (New Jersey, United States). Briefly, the next-generation sequencing library was constructed using the MetaVX Library Preparation Kit (Azenta US, Inc). For the 16s rDNA gene, the amplicons generated covered the V3 and V4 hypervariable regions with the forward primer (5’ CCTACGGRRBGCASCAGKVRVGAAT 3’) and the reverse primer (5’ GGACTACNVGGGTWTCTAATCC 3’). The 18s rDNA gene was amplified at the V4 hypervariable region with the forward primer (5’ GGCAAGTCTGGTGCCAG 3’) and reverse primer (5’ ACTACGACGGTATCTRATCRTCTTCG 3’). Next-generation sequencing was carried out on an Illumina Novaseq Platform (Illumina, San Diego, United States). Automated cluster generation and 250 bp paired-end sequencing with dual reads were performed according to the manufacturer’s instructions. The raw read files were uploaded to National Centre for Bioinformatic information (NCBI) Sequence Read Archive (SRA) with Accession No. PRJNA1093099.

The data were processed with open-source Quantitative Insights into Microbial Ecology 2 (version 2023.9) (45). The raw sequences were processed with cutadapt (46) to trim primers and adapters sequences and also remove low-quality sequences (quality threshold < 30 Phred score). The high-quality clean reads were then merged using VSEARCH (47). Sequences that failed to merge or did not contain primers were discarded. Subsequently, the reads were then processed with deblur (48) which denoised the sequences and removed chimeras. The resulting ASVs were assigned taxonomies using the SILVA 138 reference database (49). To determine the differences in community structure between the farm samples from different farms, PCoA was performed using beta-diversity metrics (weighted UniFrac, unweighted UniFrac and Bray-Curtis dissimilarity) after rarefaction to 4000 sequences per sample. Microbiome composition was analyzed by constructing taxonomic barplots at the phyla level and a heatmap at the genera level using the ggplot2 (version 3.4.4) (50) and complex heatmap (version 2.15.4) (51) packages respectively in Rstudio (version 2023.12.1).

### Biofilm former isolation and biofilm-forming evaluation

Ten colonies with diverse morphologies were randomly picked up from the PCA plates of the swab samples from the interior surfaces of the water tanks and subjected to gram staining.

The 10 bacterial isolates and an *S.* Brunei strain previously isolated from the irrigation water of a hydroponic farm (13) were cultured in tryptone soya broth (TSB; Oxoid) for 24 h at 30°C (for the 10 environment isolates) and 37°C (for *S.* Brunei), respectively. The bacterial suspensions were subsequently washed twice in PBS and diluted with a hydroponic nutrient solution to 5 Log CFU/mL and loaded to 6-well plates (2 mL/well). The hydroponic solution was formulated by mixing solution A and solution B (Watercircle Hydroponics Pte Ltd.) at 1:1 and diluted with tap water to an electrical conductivity of 2 mS/cm, the solution was then filtered with 0.22 μm filter (Millipore) for sterilization purposes. After a 5-day incubation at 25°C, each well was washed three times with sterile deionized water, stained with 2 mL 0.1% (wt/vol) crystal violet (Sigma-Aldrich) solution for 20 min at room temperature, rewashed three times with deionized water, and incubated with 2 mL of 95% ethanol at 4°C for 45 min. The absorbance reading was conducted at 595 nm using a microplate photometer (IVD model, MultiskanTM FC, Thermo Fisher Scientific Inc., MA, United States).

The biofilm-forming capacity (measured by the total biofilm biomass) was determined from the criteria established by Stepanovíc et al. (15). The optical density cut-off value (OD_c_) was the sum of the average OD of the negative control and three times the standard deviation of the negative control. Biofilm-forming ability was recognized to be “weak” if OD < 2ODc, to be “moderate” if 2ODc < OD < 4ODc, and to be “strong” if OD > 4ODc.

### WGS analysis of the strong biofilm formers C2 and C3

DNA extraction was performed using the GeneJET Genomic DNA Purification Kit (Thermo Fisher Scientific, Vilnius, Lithuania) according to the manufacturer’s instructions. The WGS was conducted at NovogeneAIT Genomics Singapore Pte Ltd. using Illumina Hiseq4000 with 350 bp insert DNA library preparation. The raw read files have been uploaded to NCBI with Accession No. SAMN40647016 and SAMN40647017. These raw reads were then assembled into contigs using SPAdes (version 3.15.3) (52). The assembled contigs were used to identify isolate species via the KmerFinder tool (version 3.2) (53) available at the Center for Genome Epidemiology (CGE) by the Technical University of Denmark (https://cge.food.dtu.dk/services/KmerFinder/).

14 *P. mexicana* and 28 *C. tuberculostearicum* genomes were downloaded from NCBI database and a phylogenetic analysis was done with the assembled contigs of the isolates. The analysis was done based on the single nucleotide polymorphism tree using CSI Phylogeny (version 1.4) (54) at CGE. The phylogenetic tree was visualized using FigTree (version 1.4.4).

### Effect of pre-formed C2 and C3 biofilms on *Salmonella* adhesion on PVC coupons

As illustrated in Figure 3a, fully grown *C. tuberculostearicum* C2 and *P. mexicana* C3 suspensions were washed with PBS and diluted with a sterile hydroponic nutrient solution prepared as described above to 5 Log CFU/mL. Each PVC coupon (2 cm × 4 cm, food-grade materials purchased from a local market in Singapore, pre-sterilised by autoclave) was submerged in 20 mL bacterial suspension and incubated at 25°C for 5 days.

After 5 days of incubation, the coupons were rinsed with sterile deionized water. Each coupon was then dipped in a 5-Log CFU/mL *S.* Brunei suspension for 5 s and rinsed thrice with sterile deionized water. Thereafter, each coupon was placed into 20 mL fresh hydroponic nutrient solution and incubated at 25°C for another 5 days. At 0 h, 24 h, 72 h, and 120 h, the bacteria were collected from the individual coupons by cell scrapers and enumerated on Xylose Lysine Deoxycholate (XLD, Oxoid) agar, a selective culture medium for *Salmonella*.

### Single– and dual-species biofilm formation by C2, C3, and *Salmonella*

Fully grown *C. tuberculostearicum* C2, *P. mexicana* C3, and *S.* Brunei cultures were washed with PBS and diluted with a sterile hydroponic nutrient solution prepared as described above to 5 Log CFU/mL. Sterile PVC coupons were submerged in 20 mL bacterial suspension (either single or mixed suspension as illustrated in Figure 3b) and incubated at 25°C for 5 days.

After 5 days of incubation, the coupons were rinsed with sterile deionized water. The bacteria were collected from the individual coupons by cell scrapers and plated on XLD (Oxoid) agar plates for *Salmonella* enumeration and on tryptone soya agar (TSA; Oxoid) plates for total bacteria enumeration.

### Efficacy of sanitizers on C2 and C3 biofilms on PVC coupons

Fully grown *C. tuberculostearicum* C2 and *P. mexicana* C3 cultures were washed with PBS and diluted with a sterile hydroponic nutrient solution prepared as described above to 5 Log CFU/mL. Sterile PVC coupons were submerged in 20 mL bacterial suspension and incubated at 25°C for 5 days.

After 5 days of incubation, the coupons were rinsed with sterile deionized water and submerged in different sanitizer solutions including 50 and 500 ppm NaOCl (Clorox® Bleach, free and total chlorine concentrations determined by sodium thiosulfate titration method), 1.5% and 3.0% H_2_O_2_, 1.0% SPC (20 mL sanitizer for each coupon in each tube). After 1 h, 6 h, and 12 h, the coupons were taken out from the sanitizer solutions and rinsed with sterile deionized water. The bacteria were collected from the individual coupons by cell scrapers and enumerated on TSA (Oxoid) plates.

### Efficacy of NaOCl and H_2_O_2_ in sanitizing NFT hydroponic systems

The sanitization trials were conducted at the same farms with the microbial analysis study between October 2023 and January 2024. All three farms had maintained continuous operation for a period exceeding three months before the sampling. After harvest, NaOCl and H_2_O_2_ were directly added to the hydroponic solutions (∼200 L hydroponic solution in each system) reaching final concentrations of 500 ppm NaOCl and 3.0% H_2_O_2_. The solutions continued to circulate in the systems with a flow rate of 0.06 m/s (same as the growing period). 6 h and 12 h later, the hydroponic solutions with sanitizers were drained and swabbing sampling and bacteria enumeration were performed from the seven spots as described above.

### Germination rate and yield of komatsuna grown in sanitized hydroponic systems

After sanitizing one hydroponic system with 500 ppm NaOCl for a duration of 12 h, the system was drained and subsequently refilled with fresh hydroponic solution. In parallel, a control system was established without any sanitization treatment for comparative purposes. To assess the impact of chemical residues on seed germination, 10 mL of the hydroponic solution from the sanitized system and 10 mL from the control system were used to germinate 100 komatsuna seeds each. The germination rate was recorded on Day 7 (n = 100). 30 seedlings from each were then randomly selected and transferred back to the two hydroponic systems (sanitized and unsanitized). 28 days later at the end of the growth circle (Day 35), the fresh weight and the number of leaves of each crop were assessed respectively (n = 30).

### The chlorination by-product analysis

At the end of the growth of komatsuna (Day 35), the hydroponic solution from the sanitized systems (without rinsing or rinsed twice with tap water, each time for 1 hour with a flow rate of 0.06m/s) was collected for analysis of a panel of chlorination by-products (Table 3a). The analysis was conducted at Eurofins Mechem Pte Ltd., according to standardized methods as listed in Table 3a. The chlorate levels in the harvested komatsuna were analyzed at SGS Testing & Control Services Singapore Pte Ltd., according to the standardized method QuPPe-PO (EURL-SRM ver. 12.2.2023) as listed in Table 3b.

### Statistical analysis

Statistical analyses were performed using Microsoft 365, Excel 2021. A paired t-test was used for data with two groups, one-way analysis of variance was used for data with more than two groups. Significant differences were considered when *P* was < 0.05.

## ACKNOWLEDGEMENTS

This study was supported by a Singapore Food Story (SFS) R&D Programme Theme 3 Competitive Grant on Food Safety and Consumer Science (W22W3D0001) funded by the Singapore Food Agency.

## AUTHOR CONTRIBUTION STATEMENT

CATH and DL conceived and designed research. CATH and GSYN conducted experiments. YHZ, MMZT and JYLT contributed analytical tools. CATH, YHZ, and MMZT analyzed data. BLP, WZ, and DL secured grants and supervised the projects. CATH and DL wrote the manuscript. All authors read and approved the manuscript.

## Figure legends

**Figure S1**. The identified bacteria genera by 16S rDNA sequencing (> 1%) from selected samples.

**Figure S2**. The identified fungi and algae phyla by 18S rDNA sequencing (> 1%) from selected samples.

## REFERENCES

1. Bloem S, de Pee S. 2017. Developing approaches to achieve adequate nutrition among urban populations requires an understanding of urban development. Global Food Security 12:80–88.

2. SFA. 2023. Food Farms in Singapore. https://www.sfa.gov.sg/food-farming/food-farms/farming-in-singapore. Retrieved 29 March 2024

3. SFA. 2022. Singapore Food Statistics 2021. https://www.sfa.gov.sg/docs/default-source/publication/sg-food-statistics/singapore-food-statistics-2021.pdf.

4. Godge M. 2022. Hydroponics: Getting to the Root of the Myths. Singapore Food agency,

5. Resh HM. 2022. Hydroponic food production: a definitive guidebook for the advanced home gardener and the commercial hydroponic grower. CRC press.

6. Ge C, Rymut S, Lee C, Lee J. 2014. *Salmonella* internalization in mung bean sprouts and pre-and postharvest intervention methods in a hydroponic system. Journal of food protection 77:752–757.

7. Li Y, Zwe YH, Tham CA, Zou Y, Li W, Li D. 2022. Fate and mitigation of *Salmonella* contaminated in lettuce (*Lactuca sativa*) seeds grown in a hydroponic system. Journal of Applied Microbiology. 132(2):1449–56.

8. Popa GL, Papa MI. Salmonella spp. infection-a continuous threat worldwide. 2021. Germs. 11(1):88.

9. WHO. 2015. WHO estimates of the global burden of foodborne diseases: Foodborne Disease Burden Epidemiology Reference Group 2007–2015. WHO, Geneva, Switzerland. https://apps.who.int/iris/handle/10665/199350.

10. Aiyedun SO, Onarinde BA, Swainson M, Dixon RA. 2021. Foodborne outbreaks of microbial infection from fresh produce in Europe and North America: a systematic review of data from this millennium. International Journal of Food Science & Technology. 56(5):2215–23.

11. Grivokostopoulos NC, Makariti IP, Tsadaris S, Skandamis PN. 2022. Impact of population density and stress adaptation on the internalization of *Salmonella* in leafy greens. Food Microbiology. 106:104053.

12. McClure M, Whitney B, Gardenhire I, Crosby A, Wellman A, Patel K, McCormic ZD, Gieraltowski L, Gollarza L, Low MS. 2023. An outbreak investigation of *Salmonella* Typhimurium illnesses in the United States linked to packaged leafy greens produced at a controlled environment agriculture indoor hydroponic operation–2021. Journal of food protection 86:100079.

13. Tham CAT, Zwe YH, Li D. 2021. Microbial study of lettuce and agriculture water used for lettuce production at Singapore urban farms. Food Control 126:108065.

14. Lipsman V, Shlakhter O, Rocha J, Segev E. 2024. Bacteria contribute exopolysaccharides to an algal-bacterial joint extracellular matrix. npj Biofilms and Microbiomes. 10(1):1–6.

15. Stepanović S, Vuković D, Hola V, Bonaventura GD, Djukić S, Ćirković I, Ruzicka F. 2007. Quantification of biofilm in microtiter plates: overview of testing conditions and practical recommendations for assessment of biofilm production by *staphylococci*. Apmis. 115(8):891–9.

16. Altonsy MO, Kurwa HA, Lauzon GJ, Amrein M, Gerber AN, Almishri W, Mydlarski PR. 2020. *Corynebacterium tuberculostearicum*, a human skin colonizer, induces the canonical nuclear factor-κB inflammatory signaling pathway in human skin cells. Immunity, Inflammation and Disease, 8(1), pp.62–79.

17. Mule P, Patil N, Gaikwad S. 2018. *Corynebacterium tuberculostearicum* a potential pathogen in breast abscess—a case report. Int. J. Med. Microbiol. Trop. Dis, 4, pp.42–44.

18. Selvaraj GK, Wang H, Zhang Y, Tian Z, Chai W, Lu H. 2022. Class 1 In-Tn5393c array contributed to antibiotic resistance of non-pathogenic *Pseudoxanthomonas mexicana* isolated from a wastewater bioreactor treating streptomycin. Science of The Total Environment, 821, p.153537.

19. Pouget C, Dunyach-Remy C, Magnan C, Pantel A, Sotto A, Lavigne J-P. 2022. Polymicrobial biofilm organization of *staphylococcus aureus* and *pseudomonas aeruginosa* in a chronic wound environment. International journal of molecular sciences 23:10761.

20. El-Azizi M, Starks S, Khardori N. 2004. Interactions of *Candida albicans* with other *Candida* spp. and bacteria in the biofilms. Journal of applied microbiology 96:1067–1073.

21. Zwe YH, Yadav M, Ten MMZ, Srinivasan M, Jobichen C, Sivaraman J, Li D. 2022. Bacterial antagonism of *Chromobacterium haemolyticum* and characterization of its putative type VI secretion system. Research in Microbiology 173:103918.

22. McGlynn W. 2004. Guidelines for the use of chlorine bleach as a sanitizer in food processing operations. Oklahoma Cooperative Extension Service.

23. Abdelshafy AM, Hu Q, Luo Z, Ban Z, Li L. 2024. Hydrogen peroxide from traditional sanitizer to promising disinfection agent in food industry. Food Reviews International. 40(2):658–90.

24. Mileto D, Mancon A, Staurenghi F, Rizzo A, Econdi S, Gismondo MR, Guidotti M. 2021 Inactivation of SARS-CoV-2 in the liquid phase: are aqueous hydrogen peroxide and sodium percarbonate efficient decontamination agents?. ACS Chemical Health & Safety. 28(4):260–7.

25. Wang R, Kalchayanand N, Schmidt JW, Harhay DM. 2013. Mixed biofilm formation by Shiga toxin–producing Escherichia coli and Salmonella enterica serovar Typhimurium enhanced bacterial resistance to sanitization due to extracellular polymeric substances. 2013. Journal of food protection. 76(9):1513–22.

26. Lin Z, Wang G, Li S, Zhou L, Yang H. Dual-species biofilms formed by Escherichia coli and Salmonella enhance chlorine tolerance. 2022. Applied and Environmental Microbiology. 88(22):e01482–22.

27. Li D, Wong CH. Isolation, characterization, and inactivation of Stenotrophomonas maltophilia from leafy green vegetables and urban agriculture systems. 2019. Frontiers in Microbiology. 10:462518.

28. Teng Z, Luo Y, Alborzi S, Zhou B, Chen L, Zhang J, Zhang B, Millner P, Wang Q. 2018. Investigation on chlorine-based sanitization under stabilized conditions in the presence of organic load. International journal of food microbiology. 266:150–7.

29. Hilgren J, Swanson KM, Diez-Gonzalez F, Cords B. Inactivation of Bacillus anthracis spores by liquid biocides in the presence of food residue. 2007. Applied and environmental microbiology. 73(20):6370–7.

30. Rodrigues PHV, Trientini M, Fisher P. 2022. Biofilm management in irrigation lines and hydroponic lettuce solutions using sanitizing chemicals. Acta Horticulturae:703–710.

31. Thakulla D. 2021. Timing and Rates of Two Hydrogen Peroxide Products to Control Algae and Evaluation of Water Temperature on Lettuce Quality for Hydroponic Production. Oklahoma State University.

32. Lau V, Mattson N. 2021. Effects of hydrogen peroxide on organically fertilized hydroponic lettuce (*Lactuca sativa* L.). Horticulturae 7:106.

33. Kriem LS, Pietzka C, Beckett M, Gärtling L, Wriedt B. 2023. Electrochemical In Situ Hydrogen Peroxide Production Can Reduce Microbial Load in Bioponic Nutrient Solutions Derived from Organic Waste. Agriculture 13:2122.

34. Thakulla D, Dunn BL, Goad C, Hu B. 2022. Timing and rates of two products using hydrogen peroxide (H_2_O_2_) to control algae in ebb and flow hydroponic systems. HortScience 57:32–39.

35. Khan MIR, Khan NA, Masood A, Per TS, Asgher M. 2016. Hydrogen peroxide alleviates nickel-inhibited photosynthetic responses through increase in use-efficiency of nitrogen and sulfur, and glutathione production in mustard. Frontiers in plant science 7:44.

36. Zhang XL, Jia XF, Yu B, Gao Y, Bai JG. 2011. Exogenous hydrogen peroxide influences antioxidant enzyme activity and lipid peroxidation in cucumber leaves at low light. Scientia Horticulturae 129:656–662.

37. Zhou J, Wang J, Shi K, Xia XJ, Zhou YH, Yu JQ. 2012. Hydrogen peroxide is involved in the cold acclimation-induced chilling tolerance of tomato plants. Plant Physiology and Biochemistry 60:141–149.

38. Goslan EH, Krasner SW, Bower M, Rocks SA, Holmes P, Levy LS, Parsons SA. 2009. A comparison of disinfection by-products found in chlorinated and chloraminated drinking waters in Scotland. Water Res. 43:4698–4706.

39. Legay C, Rodriguez MJ, Serodes JB, Levallois P. 2010. Estimation of chlorination by-products presence in drinking water in epidemiological studies on adverse reproductive outcomes: a review. Sci. Total Environ. 408:456–472.

40. Nieuwenhuijsen MJ, Toledano MB, Elliott P. 2000. Uptake of chlorination disinfection by-products; a review and a discussion of its implications for exposure assessment in epidemiological studies. J. Expo. Anal. Environ. Epidemiol. 10:586 –599.

41. Van Haute S, Sampers I, Holvoet K, Uyttendaele M. 2013. Physicochemical quality and chemical safety of chlorine as a reconditioning agent and wash water disinfectant for fresh-cut lettuce washing. Applied and environmental microbiology. 79(9):2850–61.

42. Rico D, Martin-Diana AB, Barat JM, Barry-Ryan C. 2007. Extending and measuring the quality of fresh-cut fruit and vegetables: a review. Trends Food Sci. Technol. 18:373–386.

43. Artes F, Gomez P, Aguayo E, Escalona V, Artes-Hernandez F. 2009. Sustainable sanitation techniques for keeping quality and safety of freshcut plant commodities. Postharvest Biol. Technol. 51:287–296.

44. EFSA Panel on Contaminants in the Food Chain (CONTAM). 2015. Risks for public health related to the presence of chlorate in food. EFSA Journal. 13(6):4135.

45. Bolyen E, Rideout JR, Dillon MR, Bokulich NA, Abnet CC, Al-Ghalith GA, Alexander H, Alm EJ, Arumugam M, Asnicar F, Bai Y, Bisanz JE, Bittinger K, Brejnrod A, Brislawn CJ, Brown CT, Callahan BJ, Caraballo-Rodríguez AM, Chase J, Cope EK, Da Silva R, Diener C, Dorrestein PC, Douglas GM, Durall DM, Duvallet C, Edwardson CF, Ernst M, Estaki M, Fouquier J, Gauglitz JM, Gibbons SM, Gibson DL, Gonzalez A, Gorlick K, Guo J, Hillmann B, Holmes S, Holste H, Huttenhower C, Huttley GA, Janssen S, Jarmusch AK, Jiang L, Kaehler BD, Kang KB, Keefe CR, Keim P, Kelley ST, Knights D, et al. 2019. Reproducible, interactive, scalable and extensible microbiome data science using QIIME 2. Nat Biotechnol 37:852–857.

46. Martin M. 2011. Cutadapt removes adapter sequences from high-throughput sequencing reads. EMBnet journal 17:10–12.

47. Rognes T, Flouri T, Nichols B, Quince C, Mahé F. 2016. VSEARCH: a versatile open source tool for metagenomics. PeerJ 4:e2584.

48. Amir A, McDonald D, Navas-Molina JA, Kopylova E, Morton JT, Zech Xu Z, Kightley EP, Thompson LR, Hyde ER, Gonzalez A. 2017. Deblur rapidly resolves single-nucleotide community sequence patterns. MSystems 2:10.1128/msystems.00191-16.

49. Quast C, Pruesse E, Yilmaz P, Gerken J, Schweer T, Yarza P, Peplies J, Glöckner FO. 2012. The SILVA ribosomal RNA gene database project: improved data processing and web-based tools. Nucleic Acids Res 41:D590–D596.

50. Valero-Mora PM. 2010. ggplot2: Elegant Graphics for Data Analysis. Journal of Statistical Software, Book Reviews 35:1–3.

51. Gu Z. 2022. Complex heatmap visualization. iMeta 1:e43.

52. Bankevich A, Nurk S, Antipov D, Gurevich AA, Dvorkin M, Kulikov AS, Lesin VM, Nikolenko SI, Pham S, Prjibelski AD, Pyshkin AV, Sirotkin AV, Vyahhi N, Tesler G, Alekseyev MA, Pevzner PA. 2012. SPAdes: a new genome assembly algorithm and its applications to single-cell sequencing. J Comput Biol 19:455–77.

53. Clausen PTLC, Aarestrup FM, Lund O. 2018. Rapid and precise alignment of raw reads against redundant databases with KMA. BMC Bioinformatics 19:307.

54. Kaas RS, Leekitcharoenphon P, Aarestrup FM, Lund O. 2014. Solving the Problem of Comparing Whole Bacterial Genomes across Different Sequencing Platforms. PLOS ONE 9:e104984.

